# *In situ* cryo-ET visualization of mitochondrial depolarization and mitophagic engulfment

**DOI:** 10.1101/2025.03.24.645001

**Authors:** Kevin Rose, Eric Herrmann, Eve Kakudji, Javier Lizarrondo, A. Yasemin Celebi, Florian Wilfling, Samantha C. Lewis, James H. Hurley

## Abstract

Defective mitochondrial quality control in response to loss of mitochondrial membrane polarization is implicated in Parkinson’s disease by mutations in *PINK1* and *PRKN*. Application of *in situ* cryo-electron tomography (cryo-ET) made it possible to visualize the consequences of mitochondrial depolarization at higher resolution than heretofore attainable. Parkin-expressing U2OS cells were treated with the depolarizing agents oligomycin and antimycin A (OA), subjected to cryo-FIB milling, and mitochondrial structure was characterized by *in situ* cryo-ET. Phagophores were visualized in association with mitochondrial fragments. Bridge-like lipid transporter (BLTP) densities potentially corresponding to ATG2A were seen connected to mitophagic phagophores. Mitochondria in OA-treated cells were fragmented and devoid of matrix calcium phosphate crystals. The intermembrane gap of cristae was narrowed and the intermembrane volume reduced, and some fragments were devoid of cristae. A subpopulation of ATP synthases re-localized from cristae to the inner boundary membrane (IBM) apposed to the outer membrane (OMM). The structure of the dome-shaped prohibitin complex, a dodecamer of PHB1-PHB2 dimers, was determined *in situ* by sub-tomogram averaging in untreated and treated cells and found to exist in open and closed conformations, with the closed conformation is enriched by OA treatment. These findings provide a set of native snapshots of the manifold nano-structural consequences of mitochondrial depolarization and provide a baseline for future *in situ* dissection of Parkin-dependent mitophagy.

## Introduction

Dysfunction in mitochondrial quality control in response to stress is a cellular hallmark of Parkinson’s disease (PD) ^1–4^. At the molecular level, the protein kinase PINK1 and the E3 ubiquitin ligase Parkin are implicated by human genetics in familial forms of PD ^5–7^. Mitochondrial depolarization stabilizes PINK1 on the mitochondrial outer membrane (OMM), where it phosphorylates and activates Parkin to ubiquitylate OMM proteins ^8,9^. Parkin activity leads to a host of consequences, including most famously mitophagy ^10–13^, as well as mitochondrial fission ^14^ and budding of mitochondrial-derived vesicles (MDVs)^15^.

Under basal conditions, Parkin is cytosolic, while PINK1 is constitutively degraded within the mitochondrial intermembrane space by the intramembrane protease PARL ^16–18^. Following depolarization, PINK1 import ceases, allowing for its accumulation in the translocase of the outer membrane (TOM) channel where it recruits and activates Parkin ^19,20^. PINK1 phosphorylates both Parkin and ubiquitin at Ser65 ^21^. Parkin-derived ubiquitin chains are recognized by cargo adaptors, including OPTN, NDP52, and p62 ^11–13^. Efforts by many laboratories have elucidated the structural biology of the PINK1-Parkin circuit ^22^ and of autophagy ^23^. Recently, it has become possible to structurally characterize autophagy *in situ* using cryo-FIB milling and cryo-ET ^24,25^. This raises the exciting prospect of structural visualization of the process of PINK1-Parkin-dependent mitophagy in cells. Yet PINK1-Parkin mitophagy is uniquely complex in that its upstream stimulus and depolarization has numerous and wide-ranging effects on mitochondria. Therefore, a prerequisite to understanding mitophagy at the structural level *in situ*, is to characterize the structural consequences of depolarization more broadly.

A variety of treatments are used to induce depolarization in cells. Early studies of the PINK1-Parkin pathway used the uncouplers CCCP and FCCP that non-specifically bind and transport protons to ablate the proton gradient across cell membranes ^26,27^. The most common current practice in the field is to selectively depolarize mitochondria using a combination of the F0 ATPase inhibitor Oligomycin A and the respiratory complex III inhibitor Antimycin A (OA) ^28^. Here, we established a Parkin-expressing U2OS cell line suitable for cryo-FIB milling and cryo-ET. We used this system to characterize the nano-structural consequences of OA-induced mitochondrial depolarization. We identified autophagosomes targeting depolarized mitochondria, as well as mitochondrial membrane rupture and blebbing events. Cristae became sparser and reduced in volume, and in some cases were replaced in the interior of mitochondria by vesicles. Calcium phosphate clusters completely disappeared under the OA treated condition. A subpopulation of the normally cristae-resident F0F1 ATP synthase exhibited mis-localization to the inner boundary membrane, the flat portion of the inner mitochondrial membrane (IMM) which is apposed to the OMM. We were able to reconstruct density for the intermembrane space (IMS)-resident dome-shaped prohibitin complex by sub-tomogram averaging and found that prohibitin can undergo an open-closed conformational transition, with OA favoring the closed conformation. Collectively, these data provide a nanoscale structural account of the consequences of depolarizing mitochondria.

## Results

### Validation of Parkin-mitophagy CLEM reporter U2OS cells for *in situ* cryo-ET

To visualize individual mitochondria during mitophagy initiation, we optimized Parkin-expressing U2OS cell lines for cryo-ET analysis of mitochondrial depolarization and mitophagy. We first generated a cell line that stably expressed mCherry-Parkin ^29^ as well as blue fluorescent protein targeted to the mitochondrial matrix (BFP-mito), which served as a marker for mitochondria that were competent for protein import ^30^. Consistent with previous reports in other cell lines, after treating these cells with OA for 3 h the formerly tubular and extended mitochondrial network was remodeled into spherical mitochondrial fragments ^20,29^ (Figure 1A). We observed that mCherry-Parkin was efficiently recruited to spherical mitochondria during OA treatment ^28^, while neither Oligomycin nor Antimycin A alone were sufficient to induce significant mCherry-Parkin localization to the mitochondrial surface (Figure 1A-B). To assess mitophagy flux, we generated a stable U2OS cell line that additionally expressed HALO-tagged subunit 9 of the ATP synthase (Su9-HALO ^31^), reasoning that mitochondrial damage by OA would lead to Su9 degradation. Consistently, we observed robust degradation of Su9-HALO upon OA treatment, with approximately 28% of Su9 being processed after 3 h of 3 μM OA (Figure 1C-D). Moreover, Su9 degradation was lysosome-dependent as evidenced by its reversal by the V-ATPase inhibitor Bafilomycin A1 (BafA1) (Figure 1C-D). These data show that OA-induced mitophagy proceeds in Parkin-expressing U2OS cells as expected.

**Figure 1:**
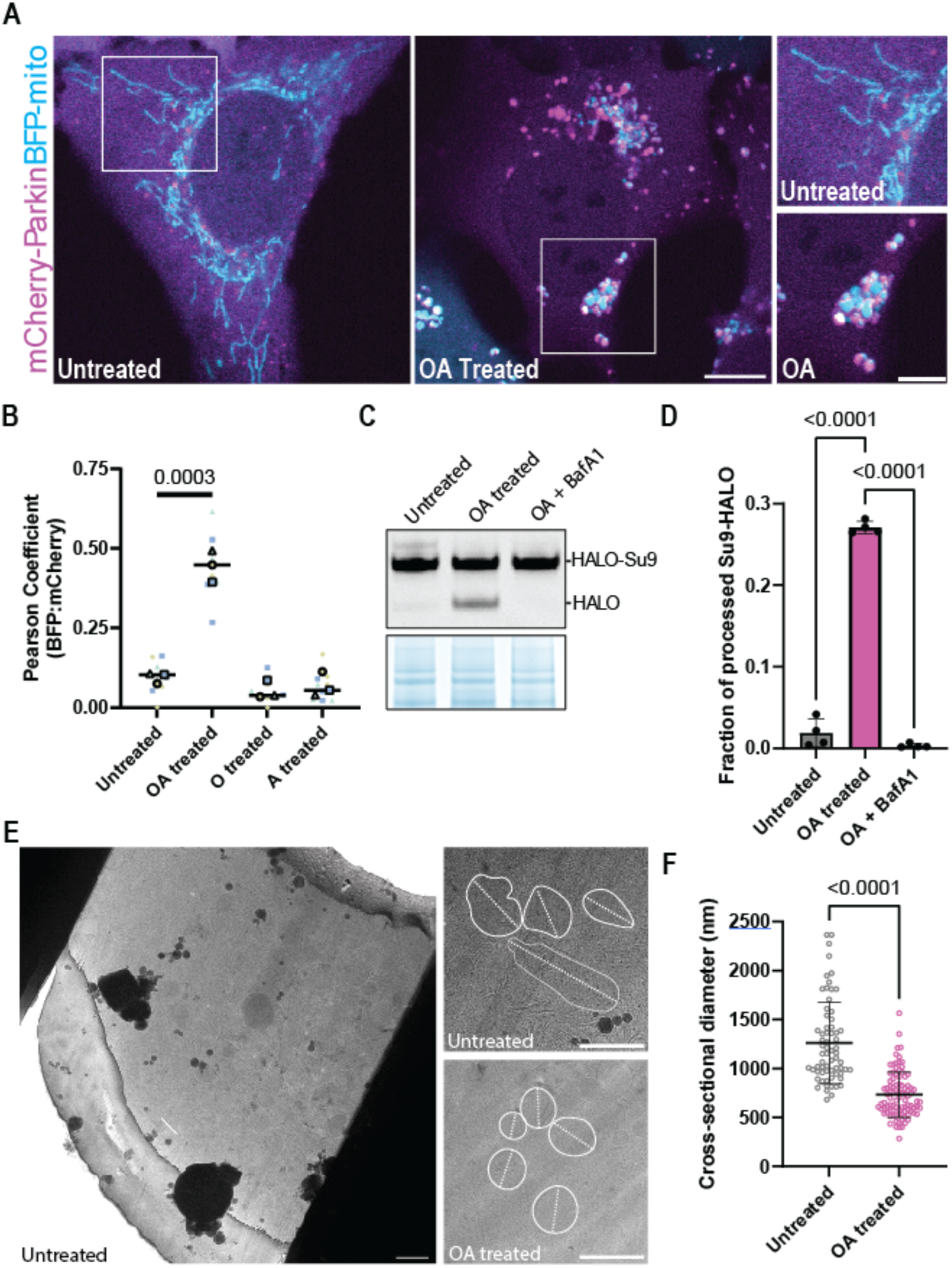
Validation of a Parkin-dependent mitophagy reporter cell line for cryo-FIB milling. Establishment and validation of CLEM mitophagy reporter cells for targeted cryo-FIB milling using mCherry-Parkin and BFP-Mito by confocal microscopy (A). Parkin recruitment to OA-induced fragmented mitochondria was only observed after dual treatment with OA but not with Oligomycin or Antimycin alone (n=3 independent fields per replicate per condition) (B). Mitophagy flux assay using the inner mitochondrial membrane protein Su9 as a probe (C). Quantification of Su9 processing in C (D). Using cryo-fluorescence of BFP-mito to guide the milling process, lamellae targeting the intact (green) or fragmented (magenta) mitochondrial network were generated (E). (F) Mitochondrial sections observable within OA-treated lamellae were significantly smaller (732 nm average diameter, n=93) than the healthy untreated network (1261 nm average diameter, n=66).

We next performed cryo-fluorescence guided cryo-FIB milling to generate lamellae containing either control or depolarized mitochondria for tomographic analysis (Fig. 1E). Upon inspection of lamellae by transmission electron microscopy (TEM), we noted mitochondrial fragmentation in OA treated, but not in untreated, cells. To further validate lamellae quality, we measured the longest cross-sectional diameter of mitochondria in these samples, which revealed a decrease in diameter from approximately 1.3 μm to 0.7 μm, consistent with depolarization-induced fragmentation (Figure 1F) ^29,32^.

In total, we generated 92 lamellae of approximately 200 nm thickness (27 untreated and 65 OA treated) for ultrastructural analysis of the mitochondrial network. We reconstructed 157 tomograms (47 untreated and 110 OA treated) and identified 141 volumes for further analysis after manual, qualitative inspection (movie 1 and movie 2 for untreated and depolarized mitochondrial tomograms, respectively). We then used Membrain to segment and isolate mitochondrial membranes in our tomograms in preparation for quantitative characterization of surface and volume measurements ^33^.

### Phagophores target and envelop mitochondrial fragments for isolation and degradation

Our membrane segmentations not only highlighted mitochondrial boundaries, but also revealed phagophores at various degrees of proximity to mitochondrial fragments following OA treatment (Figure 2A-C). Membrane segmentation analysis allowed us to make precise measurements of the curvature profile of the double membrane phagophores that targeted mitochondria. The smallest detectable phagophore was essentially flat (∼1 nm indentation). The opening of a larger phagophore grew to roughly 250 nm upon cargo engagement, and a 500 nm mitophagosome was identified completely isolating a mitochondrial fragment (Figure 2A-C insets). The ends of this phagophore cup came within 10 nm of the OMM while it was maintained at a distance of at least 16 nm from an adjacent membrane sheet. In this larger membrane-autophagosome gap, we also detected rod-like densities sandwiched between the neighboring membranes, likely ATG2A, a member of the bridge-like lipid transfer protein (BLTP) family that provides a conduit for phospholipid transfer to drive phagophore growth (Figure 2D) ^34^. Consistent with the structure of ATG2A ^35^, these densities were ∼20 nm in length, and where present, increased the membrane-autophagosome gap distance perpendicular to the membrane (movie 3). Multiple layers of putative phagophore membranes were packed tightly together, with gaps between phagophores of less than 10 nm (Figure 2E-F) (movie 4). In total, ∼17% of the mitochondrial fragments in the samples treated with OA were discernibly targeted by phagophore-like membranes (Figure 2G).

**Figure 2:**
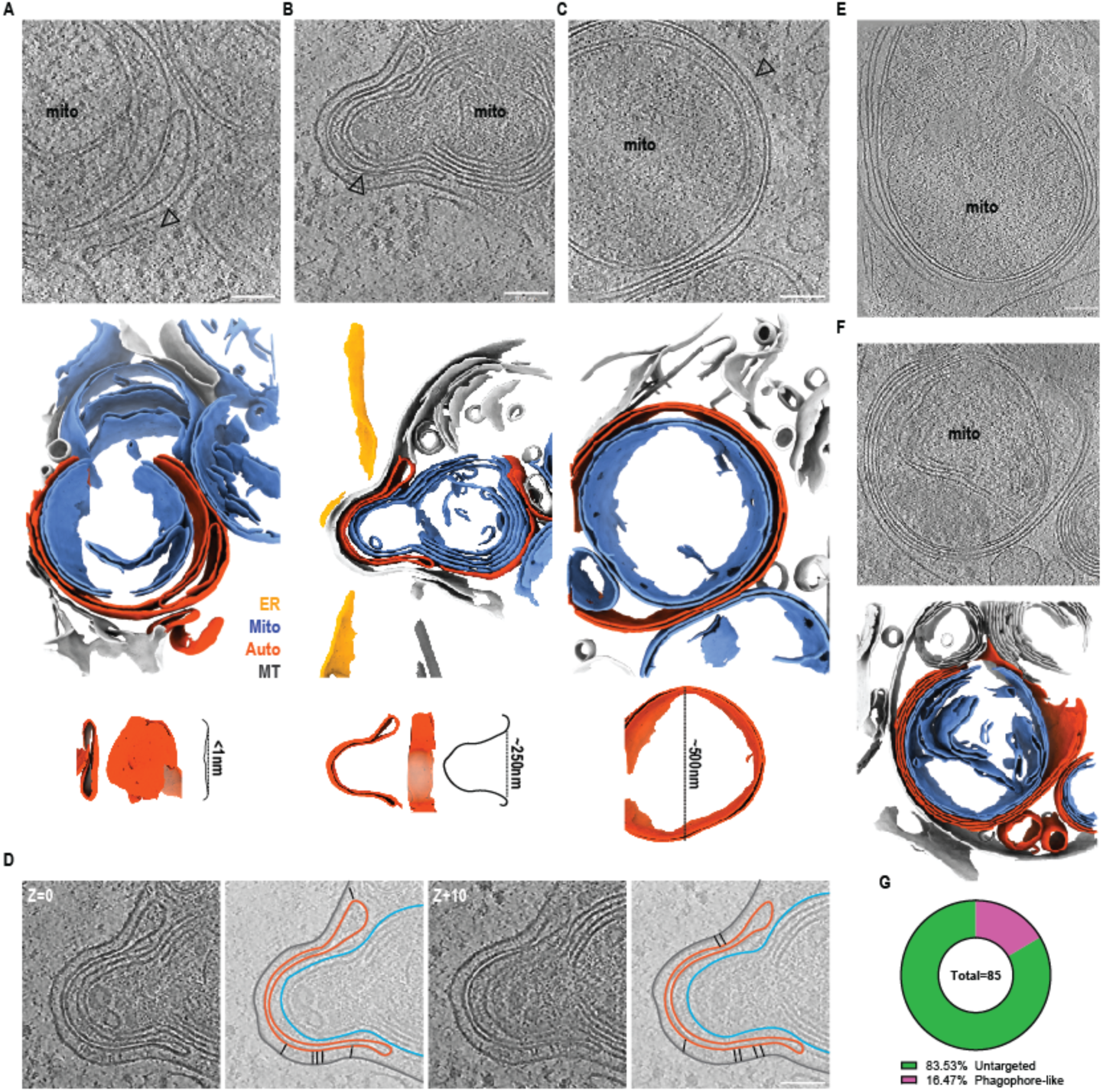
Phagophores target and envelop mitochondrial fragments. An early peanut-shaped double membrane structure that is likely an early phagophore was identified next to a mitochondrial fragment. Membrane segmentation reveals a slight dimple in this membrane structure that is less than 1nm deep (A). A larger enveloping double membrane structure with a 250nm opening was found targeting a mitochondrial fragment and adjacent to a membrane sheet (B). A fully enveloped mitochondrial fragment in a double membrane structure that is likely an early mitophagosome with a diameter of approximately 500nm (black arrows) (C). Step-wise segmentation of the volume in (B) highlighting BLTPs in between the membrane sheet and autophagosomal membranes (black sticks) (D). Two examples of mitochondrial fragments found enveloped in membrane structures consisting of more than 2 distinct membranes (E) and with membrane segmentation (F). Quantification of phagophore-like structures present in the OA treated dataset (n=85 total mitochondrial fragments, n=71 untargeted and n=14 phagophore-like) (G).

### Depolarization alters the matrix architecture of mitochondria

To investigate the nanoscale changes to mitochondria triggering mitophagy, we sought to profile ultrastructural features of mitochondria during depolarization. We observed an apparent decrease in the number of cristae per mitochondrial volume coincident with OA treatment (Figure 3A), which was substantiated by membrane segmentations (Figure 3B). Quantification of the number of cristae per mitochondrial volume revealed a statistically significant two-fold decrease in detectable segmented cristae per volume after OA treatment (Figure 3C). To generate a measure of cristae contraction with respect to the volume of the intermembrane space that they encompassed, we applied volumetric analysis of the isolated cristae, and found that the cristae surface area to volume ratio increased from 3.2 to 3.6 nm^-1^ (Figure 3D).

**Figure 3:**
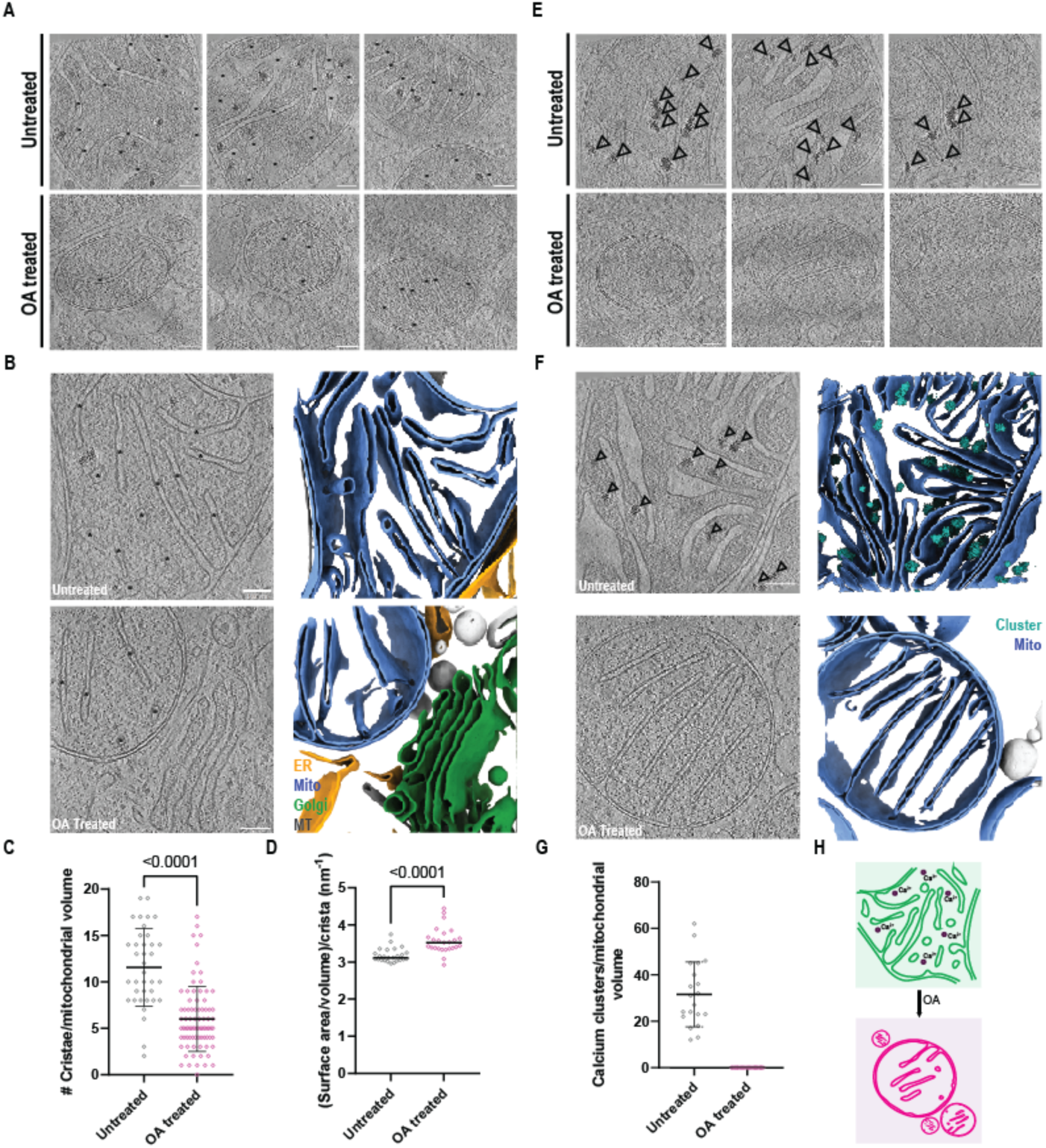
Intermembrane space shrinking and decalcification during collapse of the mitochondrial network. Cristae were abundant in untreated mitochondria and still detectable after OA treatment (black asterisks) (A). Segmentation of mitochondrial membranes illustrates cristae abundance and organization in untreated cells, with a sparser distribution following depolarization (B). (C) Quantification of membrane-segmented cristae reveals a significant decrease in average density from 12 to 6 cristae per volume after OA treatment (n=37 untreated tomograms, n=81 OA treated tomograms). (D) Cristae surface area and volume were extracted from membrane segmentations and compared between untreated and OA treated cells (n=25 independent cristae per condition, median marked). Unpaired t tests were applied to the averages from all plotted points and used to determine significance. Inspection of tomograms generated from untreated mitochondria revealed abundant matrix-resident electron dense granules, likely composed of calcium phosphate (black arrows) (E). No such clusters were detected in OA-treated mitochondrial fragments (E). Segmentation of mitochondrial membranes and calcium clusters from tomographic volumes highlights their abundance in the untreated mitochondrial network and absence in depolarized mitochondrial fragments (F). Quantification of calcium clusters reveals an average of 32 clusters were present per untreated tomogram (G). Schematic illustrating mitochondrial intermembrane space shrinking and calcium cluster loss (H).

We additionally noted abundant electron dense clusters within the mitochondrial matrix in untreated cells (Figure 3E) that were entirely absent from OA treated cells. These features were consistent with matrix granules previously reported to reside within the mitochondrial matrix ^36,37^. To characterize cluster distribution in untreated mitochondria, we back-projected a calcium cluster segmentation model trained on our dataset using Dragonfly ^38^, finding that approximately 32 clusters were found per mitochondrial tomogram volume in untreated cells, whilst zero were observed in OA treated cells (Figure 3F-H). These data show that OA treatment induces loss of matrix granules and shrinkage of the intermembrane space as indicated by sparser and thinner cristae.

### Mis-localization of ATP synthases to the IBM following depolarization

Given the alterations in mitochondrial matrix organization upon OA treatment, we next considered the structure of the inner mitochondrial membrane. The IMM comprises two morphologically and functionally distinct sub-regions, the cristae, and the inner boundary membrane (IBM). While the IBM is typically flat and in close parallel apposition to the outer membrane, the cristae are highly folded in part due to the dimerization of ATP synthase complexes that impose membrane curvature. In agreement with the known structural organization of ATP synthases in the IMM ^39^ we found numerous examples of ATP synthases forming arrays along cristae in both untreated and OA-treated cells ^40,41^ (Figure 4A). In OA treated, but only rarely in untreated, cells we identified a second class of ATP synthase molecules localized on the IBM (inset). To quantify the number of ATP synthase complexes in each class, we took the sub-tomogram average from 503 manually picked particles and trained a PyTOM model for template match-based localization ^42^ (Supplemental Figure 1A-E; Figure 4C). We first confirmed that our PyTOM picks yielded a sub-tomogram average similar to known structures of ATP synthase ^43^ (Supplemental Figure 1F), and then projected the resulting picks onto the membrane segmented tomogram and manually inspected each class (Figure 4B). Overall, we confirmed 413 cristae-associated ATP synthase complexes and 257 localized to the IBM after OA treatment, compared to 623 cristae-associated and 17 IBM-associated in untreated cells (Figure 4C). These data show that ATP synthase is mis-localized upon OA treatment, suggesting that the loss of IMM sub-domain identity is a feature of mitochondrial depolarization.

**Figure 4:**
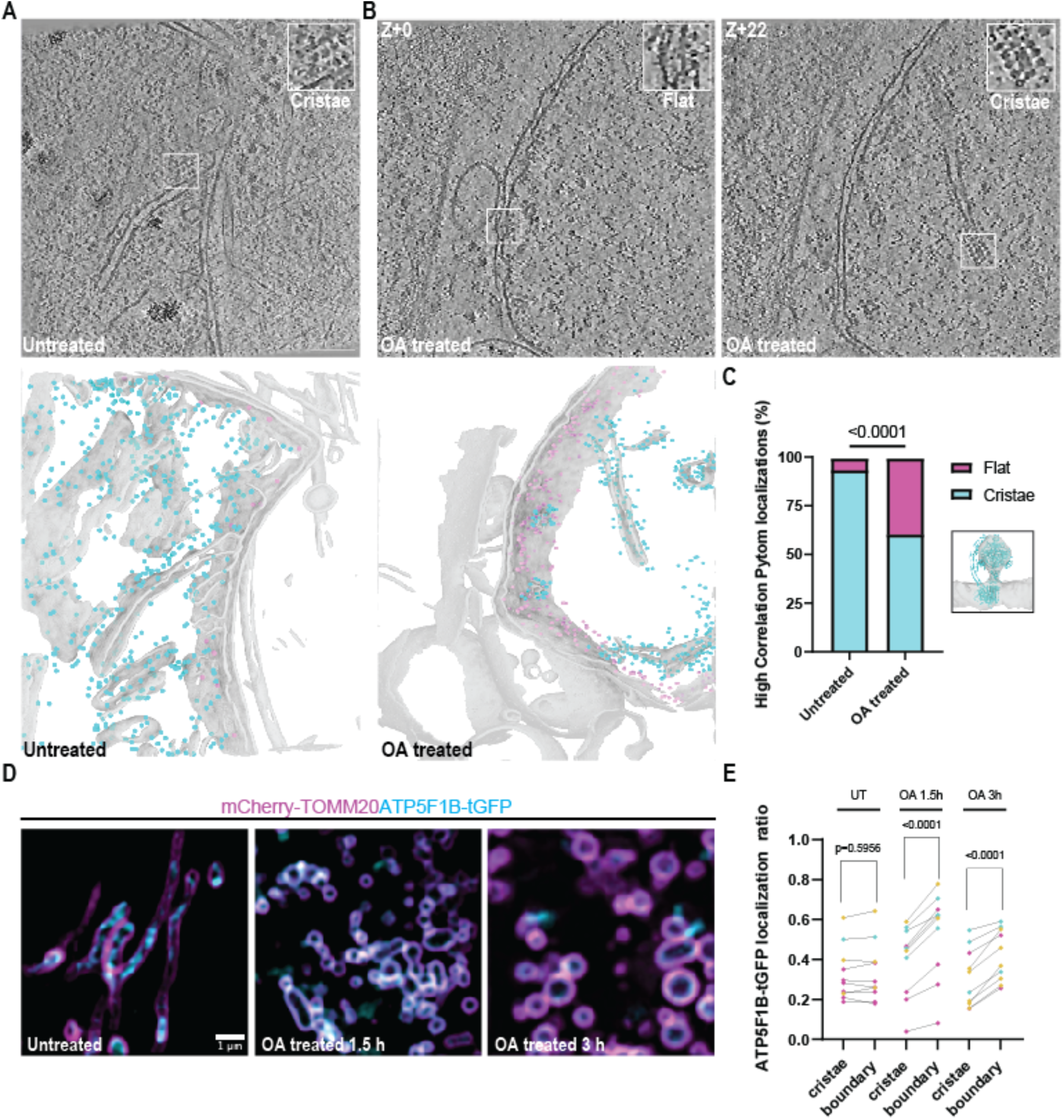
Mis-localization of ATP synthases to the inner boundary membrane after OA treatment. In untreated mitochondria, ATP synthases were found in high abundance on cristae membranes and rarely on the IBM (A and inset). After OA treatment, the OM-associated class of ATP synthases significantly increased in population (B and insets). Segmentation of the mitochondrial membrane with ATP synthase molecules identified by Pytom back projected onto cristae (cyan spheres) and the OM (magenta spheres) for visualization. Quantification of ATP synthases from each class in untreated (n=17 IBM, n=623 cristae) and OA treated cells (n=257 IBM, n=413 cristae) and comparison via Fisher’s exact test with template match density map shown and model docked (RCSB: 8H9T) (C). Live cell Airyscan imaging of mitochondria using TOMM20-mCherry and ATP5F1B-GFP to track ATP synthase localization after depolarization. (D) Quantification of localizations in (C) (n=3-4 cells per replicate over three independent replicates with a Paired t test used to determine significance).

To further characterize this mis-localization phenotype in cells actively undergoing depolarization, we employed live cell Airyscan fluorescence imaging of mitochondria for high throughput detection and analysis of ATP synthase molecules during a time course of OA treatment. We reasoned that this would permit visualization of protein localization during mitophagy initiation as early as 1.5 h after addition of OA. We transiently expressed the mitochondrial outer membrane marker mCherry-TOMM20 and ATP5F1B-turboGFP, a subunit of the human ATP synthase encoded in the nuclear genome, in U2OS cells that did not overexpress Parkin. These experiments revealed GFP-labeled cristae domains in the centers of mCherry-labeled mitochondria (Figure 4D). This central GFP density was, however, re-localized after OA treatment, as GFP signal became progressively enriched at the periphery with mCherry-TOMM20 signals. We applied 4-dimensional machine learning informed voxel segmentation to quantify the proportion of ATP5F1B-tGFP signal present in the cristae versus IBM sub-domains of the IMM, respectively, finding a statistically significant re-localization into the IBM as expected (Figure 4D). Thus, the fluorescence microscopy data confirmed the tomographic observation that ATP synthase complexes re-localize from curved cristae membranes to IBM upon OA treatment.

### Structural characterization of the Prohibitin complex and conformational changes after OA treatment

Given our observation of ATP synthase re-localization, we next considered the machinery of IMM quality control. Prohibitin complexes are dome-like structures consisting of prohibitin −1 and −2 heterodimers that assemble in the IMS via their N-terminal transmembrane helix. Prohibitin-1 and −2 are members of the SPFH protein family ^44^. Among other proposed functions, the prohibitin complex supports cristae architecture and negatively regulates the activity of the matrix-resident AAA protease (m-AAA) ^45^, which cleaves numerous proteins during import into the intermembrane space. We observed abundant prohibitin complexes in the IMS in untreated and in OA-treated cells, in contrast to the other mammalian family members stomatin and flotillin which localize to the plasma membrane and cytoplasm, respectively ^44^. Our subsequent structural determination of the prohibitin complex (see below) further supports this identification.

To determine the structure of the native prohibitin complex, we manually picked roughly 2500 prohibitin complexes from our entire dataset for subtomogram averaging (Figure 5A). 3D subclassification of the sub-tomogram averages revealed two distinct conformational states of the prohibitin complex present in untreated and OA treated cells, both of which resolved to approximately 20 Å. We termed these two conformational states “open” and “closed” on the basis of detectable protein density for the dimeric membrane-anchoring domains of the prohibitin complex (Figure 5B-C). The closed conformation manifests additional density extending 3 nm on the matrix side of the IMM, while the open complex is associated with minimal matrix density (Figure 5C).

**Figure 5:**
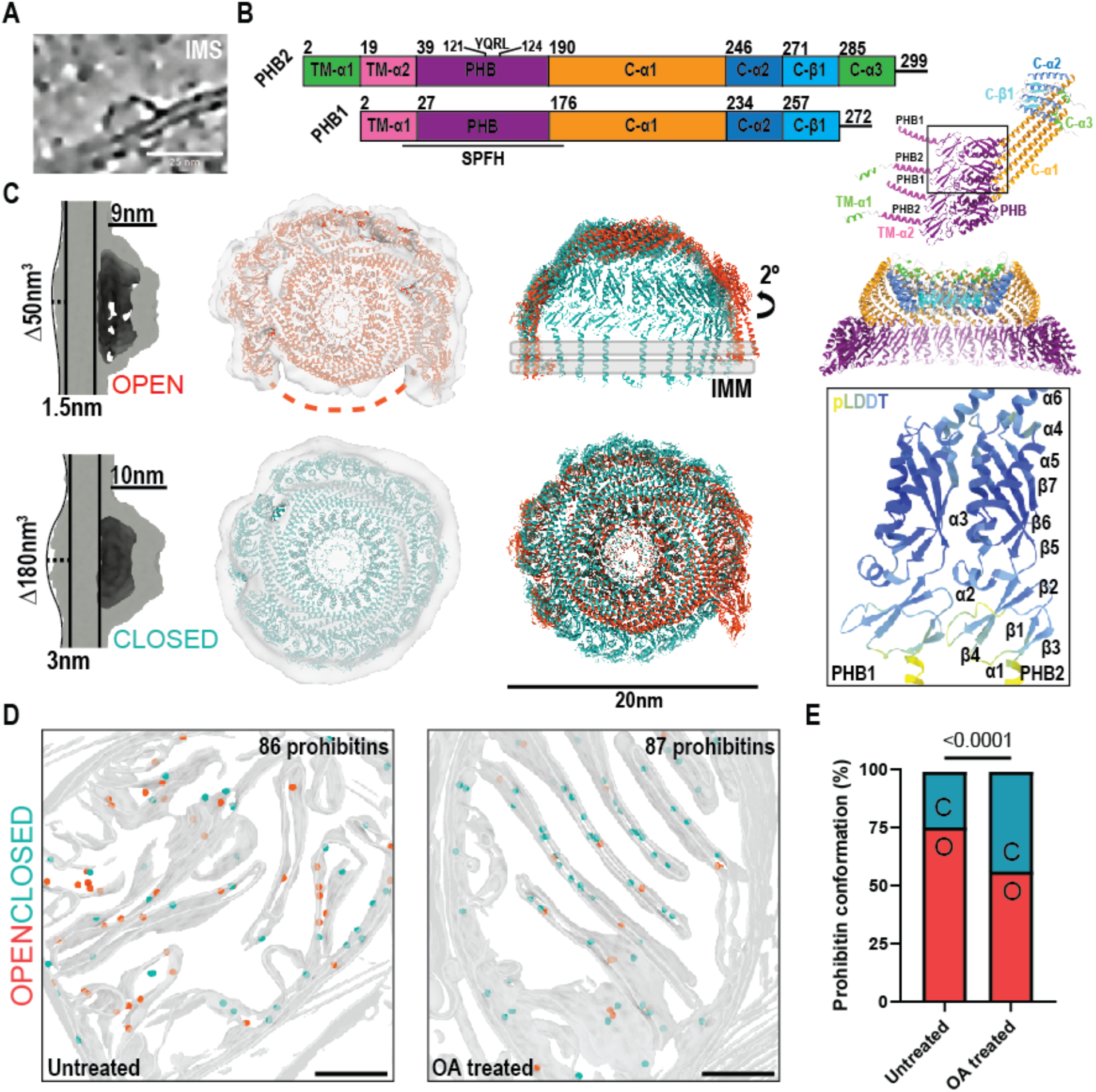
Structural determination of the prohibitin complex and its conformational transition during depolarization. Prohibitin forms a dome-like structure in the intermembrane space of mitochondria (A). Domain architecture for Prohibitin −1 and −2. A hetero-tetrameric and hetero-dodecameric AlphaFold3 predictions are also shown colored by domains, and inset shows the pLDDT scores and secondary structure architecture of the SPFH domain, annotated for Prohibitin −2 (B). (C) Side and top views of EM density maps corresponding to two solved structures of prohibitin complex highlight extra density on the matrix side of the membrane. (D) Back projection of manually picked particles (shown as spheres) onto membrane segmentations from tomograms of both untreated and OA treated mitochondria (open=salmon, closed=blue). (E) Quantification of prohibitin complexes from each class in untreated (n=930 open, n=677 closed) and OA treated cells (n=297 open, n=511 closed) and comparison via Fisher’s exact test.

Sub-tomograms processed with C1 symmetry consistently showed 12 density peaks arranged in an approximate circle. We screened 7 to 16 prohibitin 1-2 heterodimer pairs using AlphaFold3 ^46^ and found that 11-12 copies of the heterodimer pair yielded both the best pTM and iPTM scores (Supplemental Figure 2A-B). Based on the presence of 12 peaks in the density, a dodecamer of PHB1-2 dimers consisting of residues 72-272 PHB1 and 79-299 of PHB2 was generated using AlphaFold3 and automatically fit into the EM density. To improve the fit of the AlphaFold prediction, the model was iteratively relaxed into the closed then open density maps using ISOLDE ^46,47^. To generate the final full-length models including the n-terminal transmembrane helices, the N-terminus of each prohibitin was replaced by full length SPFH domains followed by a final round of ISOLDE relaxation (Figure 5C). Each prohibitin is anchored to the IMM by a transmembrane helix, with the N-terminus of prohibitin 2 folding into a second additional helix (TM-α1 and TM-α2) that may also associate with the membrane (Figure 5B-C). The SPFH domains of Prohibitin 1 and 2 pack together by contacts between helix α3 and strand β7 in individual heterodimer pairs as well as between heterodimers. The nearly 60 residue-long helix C-α1 directly continues from the SPFH domains away from the membrane and packs tightly between neighboring alpha helices from adjacent protomers. Finally, the top of the dome is assembled by tight packing of the C-terminal helix C-α2, leaving a central pore that is formed from the disordered C-terminus that forms a beta strand complementation with adjacent subunits. Prohibitin 2 contains an additional C-α3 that sticks up above the dome cap (Figure 5B). This fully assembled closed prohibitin dome is approximately 20 nm in diameter and rises 10 nm above the membrane (Figure 5C). We refer henceforward to the fully assembled state as the closed conformation. Density for all 12 heterodimer pairs was visible in our initial sub-tomogram average (Supplemental Figure 3), allowing us to model a fully enclosed dome complex. An additional unmodeled density occupying 180 nm^3^ was present on the matrix side of the membrane which cannot be accounted for by prohibitin itself. There is insufficient unmodeled mass in prohibitin 1 and 2 to account for this density, which therefore most likely reflects ordered portions of prohibitin-associated proteins or alterations to bilayer organization.

A second conformational state became evident from 3D classification of the subtomogram averages. We refer to this state as the open conformation, because density is absent for approximately 25% of the side of the wall of the dome. In the open conformation, prohibitin 1-2 heterodimers are arranged in an asymmetric spiral that are flexed outward by 2 degrees relative to the closed conformation (Figure 5C). The open complex is also roughly 20 nm in diameter and protrudes 8 nm from the IMM. In the open state, density is missing for the SPFH domains from 3 heterodimer pairs. The density quality and resolution are lower for the open state, and it seems likely that the open state density reflects an ensemble of related open states rather than a single unique conformation, as highlighted in movie 5.

To identify which complexes were in the open versus the closed conformation in our tomograms, we back-projected prohibitin complexes onto membrane segmented tomograms for analysis. In both untreated and OA treated cells, open and closed prohibitin complex conformations were identified within cristae lumens with apparent random distribution (Figure 5D). Quantification of the two populations of complexes revealed a statistically significant shift from 75% open to 56% open after depolarization (Figure 5E), indicating that the prohibitin complex is basally in a dynamic assembly of states and that depolarization drives the complex towards closure.

Back projection of Prohibitin complexes additionally allowed us to use these complexes as EM fiducials to follow large-scale disruptions in IMM morphology, even where the healthy mitochondrial ultrastructure is otherwise lost. We identified OA-treated mitochondria with spherical cristae and outer membrane peels. We found examples of prohibitin molecules exposed to the cytosol due to outer membrane peeling (Supplemental Figure 4A-B). We also found examples of single and double membrane blebs and ruptures, (Supplemental Figure 4C-E) ^48,49^ in agreement with previous studies.

## Discussion

Here, we employed cryo-ET of FIB milled lamellae from U2OS cells to characterize mitochondria after membrane depolarization. We observed wide-ranging effects at the ultrastructural and structural levels. In some cases, we revisited known consequences in greater three-dimensional and quantitative detail. In others, such as in the redistribution of F0F1 ATP synthases, and the structure of the prohibitin domain and its conformational closing upon OA treatment, the improved signal and resolution attained permitted us to discover previously unreported structural details.

Mitochondrial Ca^2+^ is implicated in normal mitochondrial function and is perturbed in neurodegenerative diseases ^50^. The soluble Ca^2+^ concentration in the mitochondrial matrix is ∼ 100 μM, much higher than in the cytosol, but lower than for other internal stores^50^. The capacity of the mitochondrion for Ca^2+^ storage is increased by the presence of solid phase calcium clusters. Here, we found abundant Ca^2+^ clusters in untreated U2OS cells, which were identified based on their similarity to previous literature reports ^36,37^. Clusters were abundant and appeared to be distributed randomly within the matrix. Depolarization of mitochondria with non-specific ionophores was previously shown to trigger Ca^2+^ cluster disappearance from the matrix ^36,37^. Here, we found that the more specific treatment with OA for OXPHOS inhibition also leads to Ca^2+^ cluster disappearance, and that at 3 h of treatment, the disappearance is complete. It is generally assumed that matrix granules consist of calcium phosphate crystals, however, it seems plausible that other phosphate-containing molecules such as nucleic acids might co-precipitate in these clusters. Their possible presence and subsequent liberation upon depolarization are significant questions for the future follow-up in the field.

We took advantage of the three dimensionality, size, and quality of the data set to quantitate changes in the cristae resulting from depolarization ^51,52^. We observed a two-fold reduction in the number of cristae after depolarization, and the remaining cristae were markedly reduced in volume. Cristae structure is known to be regulated by MICOS, OPA1, F0F1 ATP synthase, lipids, membrane potential, and calcium ^53,54^. Of these molecules, we were able to directly visualize solid-phase calcium, as described above, and F0F1 ATP synthase. In both untreated and treated cells, we were able to identify F0F1 ATP synthase molecules in the EM density and characterize their redistribution on depolarization. In untreated cells, F0F1 ATP synthases were found in ribbons on curved portions of cristae ^55,56^, as expected. The curved structure of the F0F1 ATP synthase dimer ^57,58^ is considered fundamental to stabilizing the structure of cristae. Upon OA treatment, nearly 40% of ATP synthase complexes re-localized out of the cristae and onto the IBM. This unusual localization as observed by cryo-ET was then confirmed by Airyscan fluorescence microscopy. This raises the question whether ATPase re-localization is a cause or consequence of cristae remodeling, which may be further related to the question of ATPase dimerization ^54^. The ATP synthase inhibitor oligomycin which was used in this study binds to the F0 complex at sites that could potentially influence the dimer interface ^59^. Thus, the former possibility, that ATPase inhibition by oligomycin directly influences its re-localization and cristae remodeling, seems plausible, but the latter cannot be ruled out. Since some organized F0F1 ATP synthase ribbons persist even in structurally altered cristae, ATP synthase reorganization and re-localization alone cannot fully account for cristae remodeling. Other contributors might include PINK1 regulation of MICOS via phosphorylation of its MIC60 subunit ^60^.

In this study we characterized the broad effects of mitochondrial depolarization to generate a baseline for future in depth structural analysis of Parkin-dependent mitophagy. We observed several examples of single phagophores targeting depolarized mitochondria, as well as multi-lamellar mitophagic events. Single phagophores manifested the characteristic membrane bulge at the rim as seen in other cryo-ET analyses of autophagy ^24,61^. We saw one clear example of a phagophore sandwiched between a membrane sheet and a mitochondrial fragment, potentially identified as an omegasome. Omegasomes are a specialized subdomain of the ER that contacts the nascent phagophore and provides a source of phospholipids for autophagosome growth^62^. A number of 20-nm stick-like densities were evident that spanned the gap between the omegasome and phagophore. The dimensions and arrangement of the sticks resembles that seen in a study of the bridge-like lipid transporter (BLTP) VPS13C when over-expressed in cells ^63^. The main BLTP in autophagy is ATG2A ^34,64^, and we therefore assigned the stick density in this context to ATG2A. Similar to the study of VPS13C, the endogenous ATG2A visualized here is found both near the rim region and away from it. Similar stick-like densities attributed to ATG2A were seen in an *in situ* cryo-ET study of Salmonella-phagy in HeLa cells ^24^ although these were almost entirely localized to the rim region. The observations are consistent with a “standard model” of autophagosome biogenesis ^65^ in which a high flux of phospholipid transport from the ER via ATG2A drives phagophore growth.

The multilamellar autophagic structures were reminiscent of the recently structurally characterized Salmonella-containing vacuole where multiple layers of phagophore membranes surround the pathogen ^24^. The instances of multilamellar mitophagy seen here provides a second example. This reinforces the concept that multiple rounds of initiation and expansion are sometimes needed to envelop large cargos such as bacteria and mitochondrial fragments. Most of the depolarized mitochondria not associated with single or multilamellar phagophores appeared to be only moderately abnormal, in that they contained no calcium clusters and had fewer and thinner cristae. Phagophores appeared to be associated with more severely distorted mitochondria which lacked cristae completely. We did not observe enough events to state definitively that mitophagy selectively targets the most distorted mitochondria, although the concept is conceptually appealing. Further analysis of a larger number of events will be needed to establish the structural determinants that make some mitochondrial fragments more or less preferred substrates for autophagy.

Some severely distorted mitochondrial fragments were observed that had no apparent association with phagophores. In some cases, the OMM ruptured and peeled away from the IMM, exposing prohibitin complexes in the IMM. Prohibitin has been proposed to serve as a mitophagy adaptor ^66^, however, no phagophores were detected in contact with exposed prohibitin domes. OMM blebs of ∼100 nm in diameter, which appear to correspond to budding of mitochondrial-derived vesicles (MDVs) were observed, which could represent early stage vesicles in the process of shedding for eventual degradation in the lysosome ^67^. We did not, however, visualize any direct lysosomal uptake of IMM herniations known as “VDIM”s as recently reported in several immortalized human and murine cell lines ^68^.

The Prohibitins are an evolutionarily conserved set of proteins belonging to the SPFH domain family ^44^ of proteins including the structurally well characterized members HFLK/C ^69,70^, flotillin (FLOT) ^71^ and major vault protein (MVP). Additional family members are found in prokaryotes, with the bacterial HFLC/K heterodimeric complex having undergone the most extensive structural characterization and reconstitution. Like prohibitin, the HFLC/K structure forms a closed dome-like structure that regulates the activity of its resident protease, FtsH ^69^. Recently, a new second conformational state of the HFLC/K complex bound to FtsH was solved, showing that unlike FLOT and MVP, HFLC/K can adopt both a closed nearly symmetrical dome structure as well as an open “nautilus-like” structure, likely to regulate the activity of the AAA+ protease FtsH ^70^ (Supplemental Figure 5A-B).

The structure shown here was determined for endogenous protein and symmetry was imposed. The resulting C1-symmetric structure was found to contain twelve PHB1-2 dimers. During data processing in this study, test processing in symmetries ranging from C2 to C16 improved neither the FSC nor the quality of the density, and therefore analysis was carried out in the context of the C1 reconstruction. Here, we found that like HFLC/K, prohibitin adopts both a closed nearly symmetric structure as well as an open asymmetric conformation ^69,70^. Our model fits well into a density map putatively assigned to endogenous prohibitin in Chlamydomonas ^72^ (Supplemental Figure 5D), consistent with the assignment of the density to prohibitin, and highlighting a high degree of structural conservation across biology. Common features in the HFLC/K and our C1 prohibitin structures, include extensive inter-SPFH domain contacts, a helical barrel ^71^, and a 12-stranded β-sheet formed at the tip of the dome by one C-terminal strand per subunit.

Our structures contrast with a recently reported model based on an *in situ* cryo-ET reconstruction of prohibitin in human cells in which C11 symmetry was imposed ^73^ (RCSB 8RRH, Supplemental Figure 5C). The 8RRH atomic model based on the C11-symmetrized reconstruction left large regions of the density unmodeled, and contains large gaps between subunits that are atypical of stable protein complexes. The C11 coordinates lack key common features of SPFH assemblies that are present in high resolution experimental structures of flotillin and HFLC/K (e.g. RCSB 9CZ2, Supplemental Figure 5A). The C1 reconstruction reported here fills the density, manifests inter-subunit packing typical of stable complexes, and conforms to the patterns that are by now expected in SPFH family members. Prohibitin is known to interact with and regulate the activity of the m-AAA protease in the intermembrane space to stimulate protein translocation. On the basis of its two distinct conformational states, we propose that in untreated cells where prohibitin complexes are more open in confirmation, the open state is “active” and allows access of the m-AAA to its import substrates for processing. Under depolarization, prohibitin shifts to the closed “inactive” state, which could, in principle, reduce the degradative capacity of m-AAA with respect to integral membrane proteins of cristae.

In conclusion, we generated and analyzed a large repertoire of human mitochondrial volumes undergoing depolarization and mitophagy. In doing so, we created a baseline to launch future investigations of the structural basis of PINK1/Parkin-dependent mitophagy. Many of the observations here could be fruitfully expanded upon in future work. The precise mechanism driving ATP synthase re-localization to the IBD and its contribution to cristae remodeling was not resolved. This might require sub-nanometer resolution or 3D template matching of various conformations of the complex to fully explain ^74^. Contributions of other factors to cristae remodeling, such as OPA1 ^75^, remain to further explored by in situ methods in response to depolarization. While we noticed a tendency for phagophores to be associated preferentially with more severely distorted mitochondria, the sample size collected here was insufficient to draw firm conclusions. It will be important to localize PINK1-Parkin-TOMM20 complexes ^76^ *in situ* in future work and to relate these to mitophagic uptake. A finer sampling of timepoints between the onset of autophagy initiation and mitochondrial degradation, coupled with the structural mapping of these additional components, will be needed to build on the baseline described here and so definitively reveal the molecular mechanisms of PINK1/Parkin mitophagy *in situ*.

## Supporting information

key resource table

## Acknowledgements

We thank M. Lazarou and other members of ASAP team mito911 for helpful discussions and advice, L.-M. Joubert for cryo-ET sample preparation support, and D. Tudorica for helpful advice on the mitophagy assays. A portion of this research was supported by NIH grant U24 GM139168 and performed at the Midwest Center for Cryo-ET (MCCET) and the Cryo-EM Research Center in the Department of Biochemistry at the University of Wisconsin-Madison. This research was funded by Aligning Science Across Parkinson’s [ASAP-000350] through the Michael J. Fox Foundation for Parkinson’s Research (MJFF) (J.H.H, F.W., and S.C.L.), and by the Alexander von Humboldt Foundation (E.H. and J.H.H.). Some of this work was performed at the Stanford-SLAC CryoET Specimen Preparation Center (SCSC), which is supported by the National Institutes of Health Common Fund’s Transformative High Resolution Cryoelectron Microscopy program (U24GM139166).

## Competing interests

J.H.H. is a cofounder of Casma Therapeutics and receives research funding from Hoffmann-La Roche. The other authors declare that they have no competing interests.

## Data and Materials Availability

Models, maps, and raw cryo-ET data are being deposited at the protein data bank at the RCSB, the Electron Microscopy Data Bank (EMDB), and the Electron Microscopy Public Image Archive (EMPIAR), respectively.

## Materials and Methods

### Cell culture and cell line generation

Human Osteosarcoma and HEK293T cells were received from the UCB Cell Culture Facility. Cells were cultured for no more than 20 passages in DMEM supplemented with 10% fetal bovine serum, Pen/Strep (Life Technologies, catalog number 15140122), and l-glutamine (Life Technologies, catalog number 25030081). Cells were maintained in a copper-lined Heracell VIOS 160i tissue culture incubator (ThermoFischer catalog: 51033574) at 37 °C and 5% CO2 and checked for mycoplasma contamination.

To generate mCherry-Parkin and BFP-mito lentiviruses for cell line generation 4E6 HEK293T cells were seeded into 10 cm plates and transfected the next day with 45 μL Mirus LT1 transfection reagent (MIR2300) added to a mixture of 5 μg each (15 μg total) of plasmids VSV-G (addgene: 8454), R8.74 (addgene: 22036) or CMV-Gagpol (addgene: 35614), and one of the following: pLV-mCherry-Parkin (this study), pLV-BFP-mito (this study), and Su9-GFP-HALO (addgene: 184905) in 1.5 mL Optimem (ThermoFischer catalog: 31985062) according to manufacturer recommendations. Both lentivirus plasmids were restriction subcloned using NheI and BsrGI into pMK1253 (addgene: 133058) from pBMN-mCherry-Parkin (addgene: 59419, PCR amplified to include a c-terminal BsrGI cut-site) and EBFP2-mito-7 (addgene: 55248). Plasmid sequences were confirmed by nanopore full plasmid sequencing. Supernatant containing viruses was obtained 3 days post-transfection, clarified by centrifugation at 2000 rpm for 2 minutes, and concentrated 10-fold using Lenti-X concentrator (Takara Bio catalog: 631231). On the day prior to transduction, U2OS cells were seeded at a density of 100,000 cells per well into individual wells of a 12 well plate (catalog number: 07–200-82). Stable pools of cells expressing desired proteins of interest were obtained by titrating virus concentrate to achieve near 100% expression efficiency, and passaging cells once prior to experiments. Detailed protocols may be found here: dx.doi.org/10.17504/protocols.io.81wgbxq2qlpk/v1; dx.doi.org/10.17504/protocols.io.yxmvm3z5bl3p/v1

For live time lapse imaging, U2OS cells were plated on glass-bottom 35 mm dishes (Mattek, P35GC-1.5-14-C) 24 h prior to transient plasmid transfection, and 48 h prior to imaging. Plasmid transfection occurred in Opti-MEM™ I Reduced Serum Medium (Thermo Fisher Scientific, REF: 31985-070) with Lipofectamine 2000 reagent (Thermo Fisher Scientific, REF:11668030). mCherry-TOMM20-N-10 was a gift from Michael Davidson (addgene: 55146; RRID:Addgene_55146). ATP5F1B-turboGFP was generated via custom synthesis by OriGene, based on NCBI mRNA sequence identifier NM_001686 (SKU: RG201638). Single-copy plasmid inserts were verified by Sanger sequencing. A detailed protocol may be found at dx.doi.org/10.17504/protocols.io.ewov1dr82vr2/v1.

### OA treatment and Quantification of Parkin recruitment to mitochondria

Stable U2OS cells expressing mCherry-Parkin and BFP mito were seeded into 8 chamber glass bottom plates (Fischer Scientific catalog: NC1273035) at 25,000 cells per well in 250 uL of DMEM and cultured overnight. On the proceeding day, cells were treated with fresh media for 30 minutes prior to being subjected to fresh media or mixtures of Oligomycin (Sigma catalog: SIAL-O4876-5MG) and Antimycin A (Sigma catalog: SIAL-A8674-25MG) or either compound individually, for 3 hours. After incubation, cells were immediately imaged using a using a Nikon A1 confocal microscope with a 63× Plan Apochromat 1.4 numerical aperture objective. Identical imaging settings were used across all replicates and 3 independent fields of view were captured for each condition per replicate. To quantify pearson correlations for Parkin and BFP-mito under each condition, fields of cells were analyzed in ImageJ (https://imagej.net/). Fields were first subject to background subtraction using a rolling ball radius of 50 and then binarized before using the automatic thresholding from JACoP (https://imagej.net/plugins/jacop) to determine pearson coefficients. Statistical significance was determined using an unpaired t-test in GraphPad Prism 10 to compare the triplicate averages from each experiment. A detailed protocol can be found here: dx.doi.org/10.17504/protocols.io.5qpvoox4dv4o/v1

### Mitophagy flux assay via in-gel fluorescence

Stable U2OS cells expressing mCherry-Parkin and Su9-HALO were seeded at 250,000 cells per well in a 12 well plate and incubated overnight. On the following day, cells were given fresh media for 30 minutes containing 100 nM Janelia Fluor 646 HALO ligand (Promega catalog: GA1120) prior to the addition of fresh media, OA-containing fresh media, or fresh media containing both BafA1 (Medchem catalog: HY-100558) and OA for 3 hours. Cells were then harvested via scraping, pelleted at 2500 rpm for 2.5 minutes, and lysed on ice for an hour in buffer containing: 50 mM Tris, pH 7.4, 150 mM NaCl, 1mM EDTA, 0.5% NP-40, and protease inhibitor (ThermoFisher catalog: A32963). Lysates were spun at max speed for 10 minutes and subject to protein quantification using Bio-Rad Protein Assay Reagent (Bio-Rad catalog: 5000006EDU) and a BSA standard curve. 20 μg of protein was loaded in each well. Gels were imaged using the ChemiDoc fluorescent imager (Bio-Rad catalog: 12003153) and quantified in Fiji. Briefly, individual bands corresponding to processed or unprocessed HALO were quantified by measuring all pixels within a rectangular area for each sample. The processed fraction was divided by the total number of pixels in each lane and this fraction was used to determine significance from quadruplicate replicates using an unpaired t-test in GraphPad Prism 10. A detailed protocol can be found at dx.doi.org/10.17504/protocols.io.x54v9pwkmg3e/v1.

### EM grid seeding and cryo-FIB milling

Gold quantifoil R2-2 200 mesh EM grids (EMS catalog: Q250-AR2) were glow discharged for 30 seconds at 25mA. Grids were then floated on drops of 0.01% poly-l-lysine (Sigma catalog: A-005-M) in a laminar flow hood for 30 minutes. 8 chamber slides with removable wells (Sigma catalog: PEZGS0816) were simultaneously coated with 250 μL of 0.01% poly-l-lysine. During incubation, U2OS reporter cells were split using 0.25% trypsin (Fisher Scientific catalog: 25-200-056). Poly-l-lysine was removed from the wells and replaced with fresh DMEM, and EM grids were rinsed in DMEM prior to insertion into the bottom of the well of the 8 chamber plate. Cells were resuspended to 100,000 cells/mL and 200 uL of cells were transferred to each well atop a single EM grid, such that 20,000 cells/grid is achieved. Cells on grids were left to recover overnight. On the following day cells were treated with OA media as above. During OA treatment, a vitrobot Mark IV (ThermoFisher) was equilibrated to 90% humidity and 37C temperature and blotting paper and a Teflon sheet were inserted into the chamber. After OA treatment, the removable wells were removed from the 8 chamber slide and grids were retrieved using vitrobot tweezers. Grids were washed 3 times with drops of PBS and double blotted using a blot force of 10 for 8 seconds before plunge freezing into liquid ethane. Grids were then clipped using notched grid bases for cryo-FIB milling (ThermoFisher). A detailed protocol can be found here: dx.doi.org/10.17504/protocols.io.dm6gpde65gzp/v1

Notched base-clipped EM grids were loaded into an Aquilos 2 with integrated fluorescence (iFLM, ThermoFisher). Grids were first screened using SEM to identify potential lamellae sites. Cells were then screened iteratively using iFLM to target specific regions of clustering mCherry or BFP signal for lamellae site placement. Grids were then sputter coated for 15 seconds using 30mA current and 10Pa pressure, and subsequently GIS coated for 1 minute. Auto-TEM was then used to generate a lamellae of approximately 200 nm thickness at each site. Briefly, 0.3-0.5 nA of current was used to ablate cell material to 3 μm in thickness. Then 100 pA current thinned cells to 1 μm. 50 pA was used to thin cells to 500 nm, and 30 pA was used to thin lamellae to their final 200 nm thickness.

### Cryo-electron tomography data acquisition

Grids containing lamellae were retrieved from the Aquilos and immediately stored in nitrogen or loaded into a 300 kV Titan G2, G3, or G4 Krios. Untreated cells were analyzed on the G4 equipped with a cold field emission gun (CFEG), a Selectris X Energy Filter, and a Falcon 4i direct electron detector (Thermo Fisher Scientific, Hillsboro, OR, USA). The images were acquired in EFTEM mode with a 10eV slit width. OA treated cells were analyzed on a Titan G2 or G3 using a Quantum K3 direct electron detector (Gatan) and images acquired in EFTEM mode with a 25 eV slit width. The autogrids containing lamellae were loaded such that the pre-tilt axis induced by FIB milling was perpendicular to the tilt axis of the microscope. Montage maps were generated for the entire autogrid to identify lamellae positions and a second medium mag montage generated at each lamellae site. Polygon montages were used to outline the borders of each lamellae and used to guide data collection. These polygon montages were used to make diameter measurements of mitochondria across datasets. Tilt-series were collected using a dose-symmetric scheme starting from 10-15 degree lamellae pre-tilt with increments of 3 degrees in groups of 2 tilts ^77^. The nominal defocus was varied between tilt-series from - 2 to −6 μm with a step size of 0.25 μm. The total dose per tilt series was approximately 120 e^-^/ Å^2^. Frames were saved in Electron Event Representation (EER) format for G4 data.

### Cryo-electron tomography data processing and model building

Scipion was used to facilitate all downstream data processing ^78^. Briefly, MotionCor3 was used to motion correct tilt series and binning in Fourier space to the physical pixel resolution was applied during correction ^79^. CTFfind 5 was used for CTF estimation and AreTomo2 was used for tilt series alignment and tomogram reconstruction ^80,81^. Tomograms were denoised using 3DEM and segmented using Membrain with default parameters ^33,78^, and Dragonfly models were trained as previously described ^38^. Cristae surface area and volume measurements were made using Measure and Color Blobs in ChimeraX ^82^. Significance for these measurements was determined using an unpaired t-test in GraphPad Prism 10.

For sub-tomogram averaging and template match picking, ATP synthase and prohibitin complexes were manually picked using Napari (https://www.napari-hub.org/) and imported into Relion 5 for particle extraction and downstream processing ^83^. First, an initial model was generated and then subjected to 3D refinement. The resulting average for ATP synthase was then used as a template for PyTOM. Template match picking was performed with a CC score of 0.44 and a masked search using membrane segmentations that were expanded to 100 Å to confine the search for templates. For prohibitin, particles underwent 3D classification which revealed two distinct conformations, as well as a junk class that was discarded. Particles for the two prohibitin conformation classes were reconstructed at bin1 and subjected to Bayesian polishing and CTF Refinement before another round of 3D refinement. Postprocessing of each class yielded EM density maps of approximately 20Å. To build models that fit the EM density maps for each class, AlphaFold3 was used to generate an initial protein model of 12 prohibitin 1-2 dimers which was then relaxed into either EM density map using ISOLDE ^47^. Significant differences for ATP synthase localization and prohibitin conformational changes were determined using Fisher’s Exact test in GraphPad Prism 10. A detailed protocol covering all tomography data collection to model building can be found here: dx.doi.org/10.17504/protocols.io.36wgq6rkklk5/v1

### Live Airyscan microscopy of mitochondrial ultrastructure

Live cell images were acquired using the Zeiss LSM 980 with Airyscan 2 confocal microscope using an inverted 63X, 1.4 numerical aperture oil objective. Cells were imaged in a 37 °C humidified chamber with 5% CO2. Arivis Vision4D (ver 4.2.1) was used for image analysis. The z-stacks images were processed using 3D Airyscan processing from the Zeiss ZEN Blue software version 3.7 (Carl Zeiss) and saved as 16-bit czi files. The czi files were converted to the Arivis sis format for analysis.

ATP5F1B-tGFP was segmented using the Watershed option. Watershed was first performed on individual images and the optimal threshold was defined as allowing sufficient segmentation of the ATP5F1B-tGFP signal without over-segmenting on a per image basis. Then, the threshold of each image was divided by the intensity of ATP5F1B-tGFP captured in the given image resulting in a normalization factor. Finally, the median of the normalization factors was used to give us a final normalization factor of 0.26. The intensity of ATP5F1B-tGFP of each image was multiplied to 0.26 to allow a standardized threshold across all images. Segmentation accuracy was confirmed by manual review.

Segmentation of mCherry-TOMM20-N-10 and of the regions enclosed by mCherry-TOMM20-N-10 was done using Arivis’ machine learning segmenter. The segmenter was trained using 9 representative images, 3 per condition, classifying the mCherry-TOMM20-N-10, regions enclosed by mCherry-TOMM20-N-10, and background until adequate segmentation was achieved and artifacts kept to a minimum.

Intersection between the ATP5F1B-tGFP segments acquired via Watershed and either the mCherry-TOMM20-N-10 segments or the segments ROIs of the regions enclosed by mCherry-TOMM20-N-10 were performed using the “Object math” option. The surface area of the ATP5F1B-tGFP segments intersecting with the mCherry-TOMM20-N-10 segments or with the regions enclosed by mCherry-TOMM20-N-10 segments were defined as “ATP5F1B in the boundary domain” or “ATP5F1B in the cristae domain”, respectively. Each value of ATP5F1B in the boundary domain and ATP5F1B in the cristae domain was normalized to the surface area of the mCherry-TOMM20-N-10 segments ROIs and represented in graphs. Statistical analyses were performed in GraphPad Prism (ver 10.1.0), using the paired Student’s t-test with a two-tailed p-valued (p<0.05).

---

**Supplemental Figure 1:**
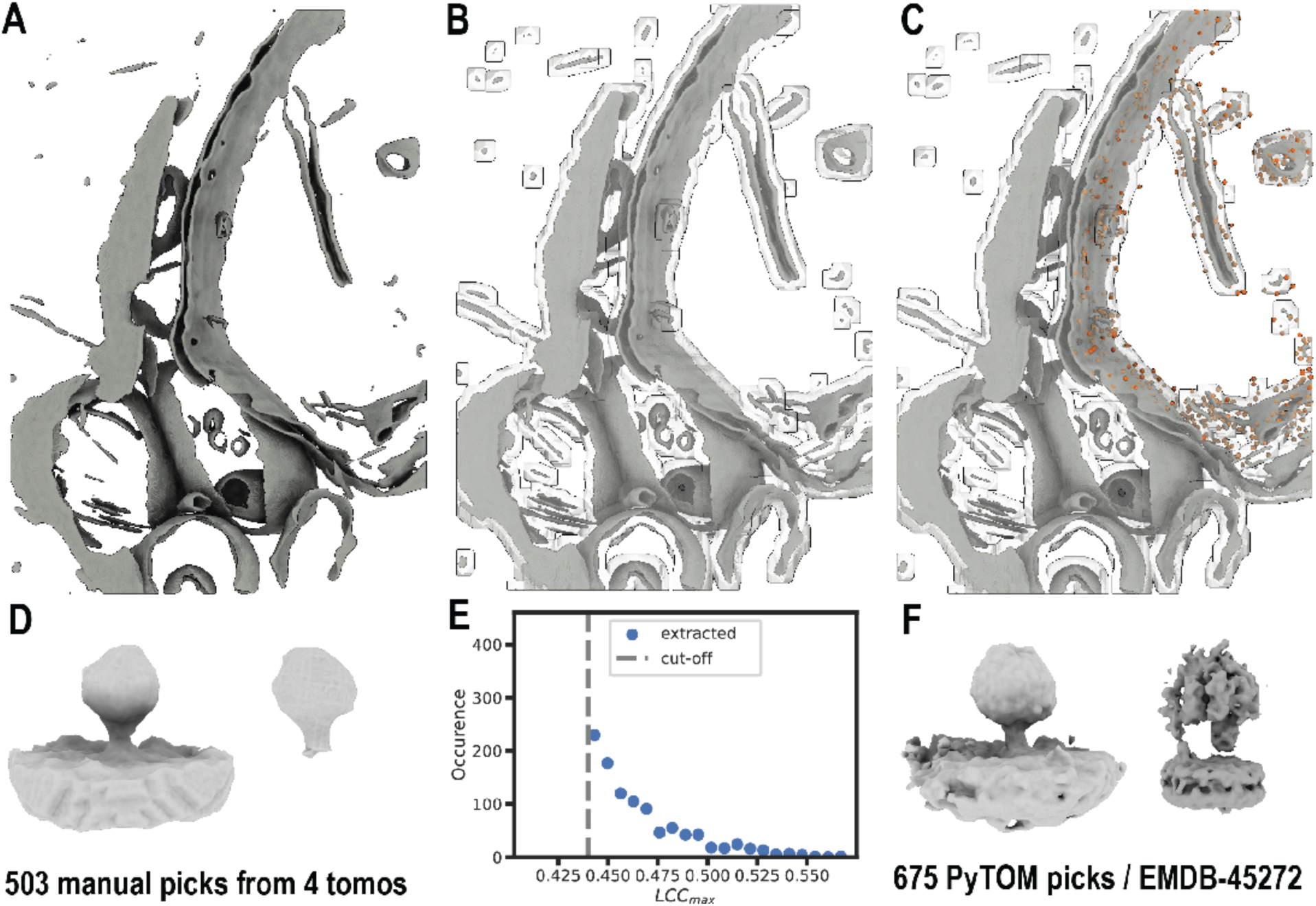
Template match picking strategy for ATP synthase. Membranes from tomograms were first segmented to guide the particle picking process (A). A mask to guide template match picking was generated using the membrane segmentation (B). Output of template match picking (C) using the resulting sub-tomogram average from 503 manual picks of 4 tomograms with membrane (left) or without (right) (D). (E) PyTOM parameters used to restrain template matching. Initial sub-tomogram average from 675 template match picks from PyTOM compared to a recently solved density map (EMDB-45272) (F).

**Supplementary Figure 2:**
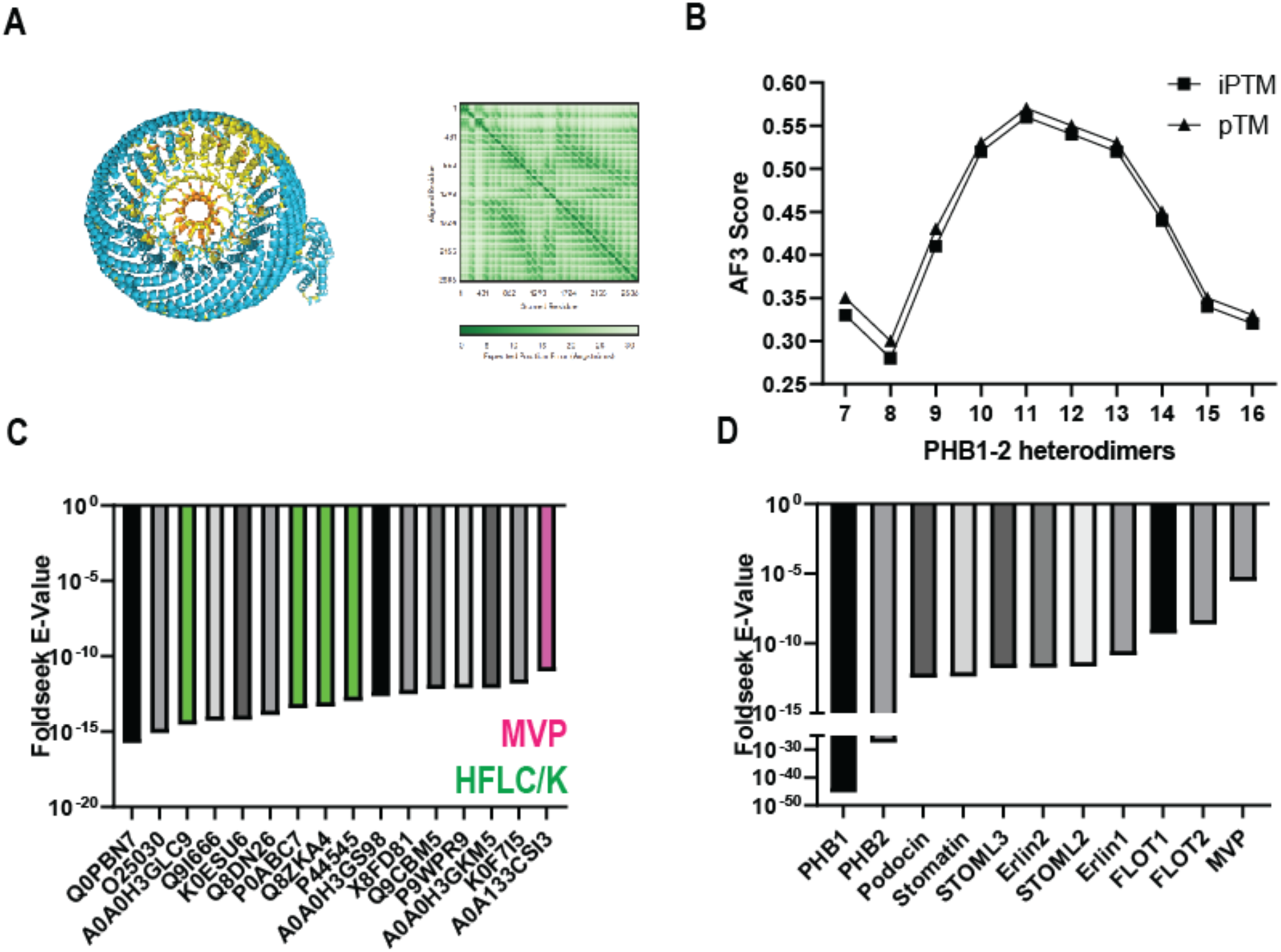
Structural analysis and comparison of Prohibitin and the closely related HFLK/C complexes. Alphafold modeling of a heterododecamer Prohibitin 1-2 complex and resultant pLDDT plot from Alphafold 3 with 2 heterodimer copies (A). Alphafold 3 screen using 2 heterodimer copies shows a preferred stoichiometry between 11 and 12 heterodimer copies (B). Foldseek using the structure of human prohibitin shows greater E-Values to bacterial HFLC/K structures than to other closely related mammalian SPFH domain-containing proteins (C-D).

**Supplementary Figure 3:**
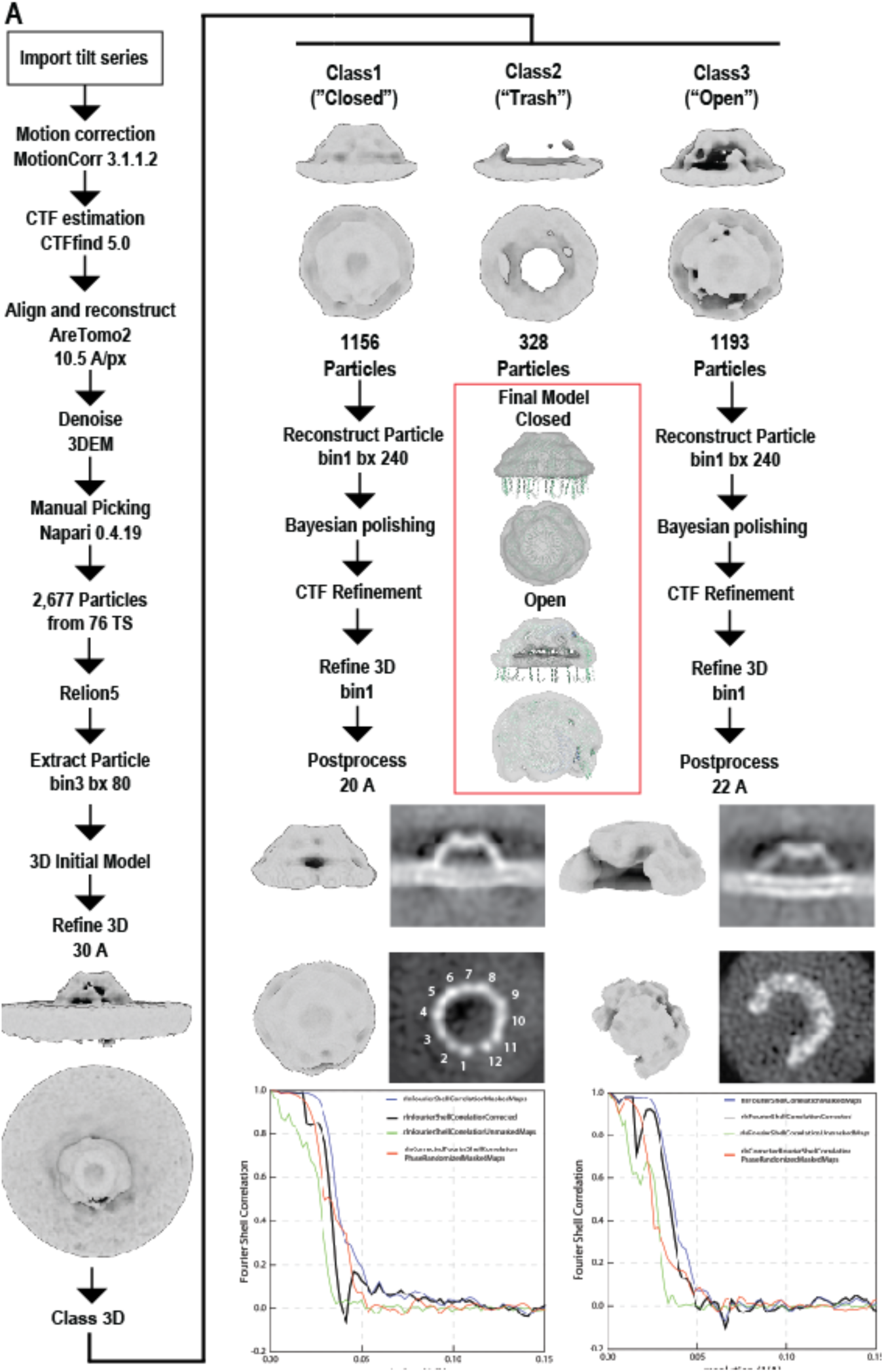
**Data processing pipeline for sub-tomogram averaging of the prohibitins complex**

**Supplemental Figure 4:**
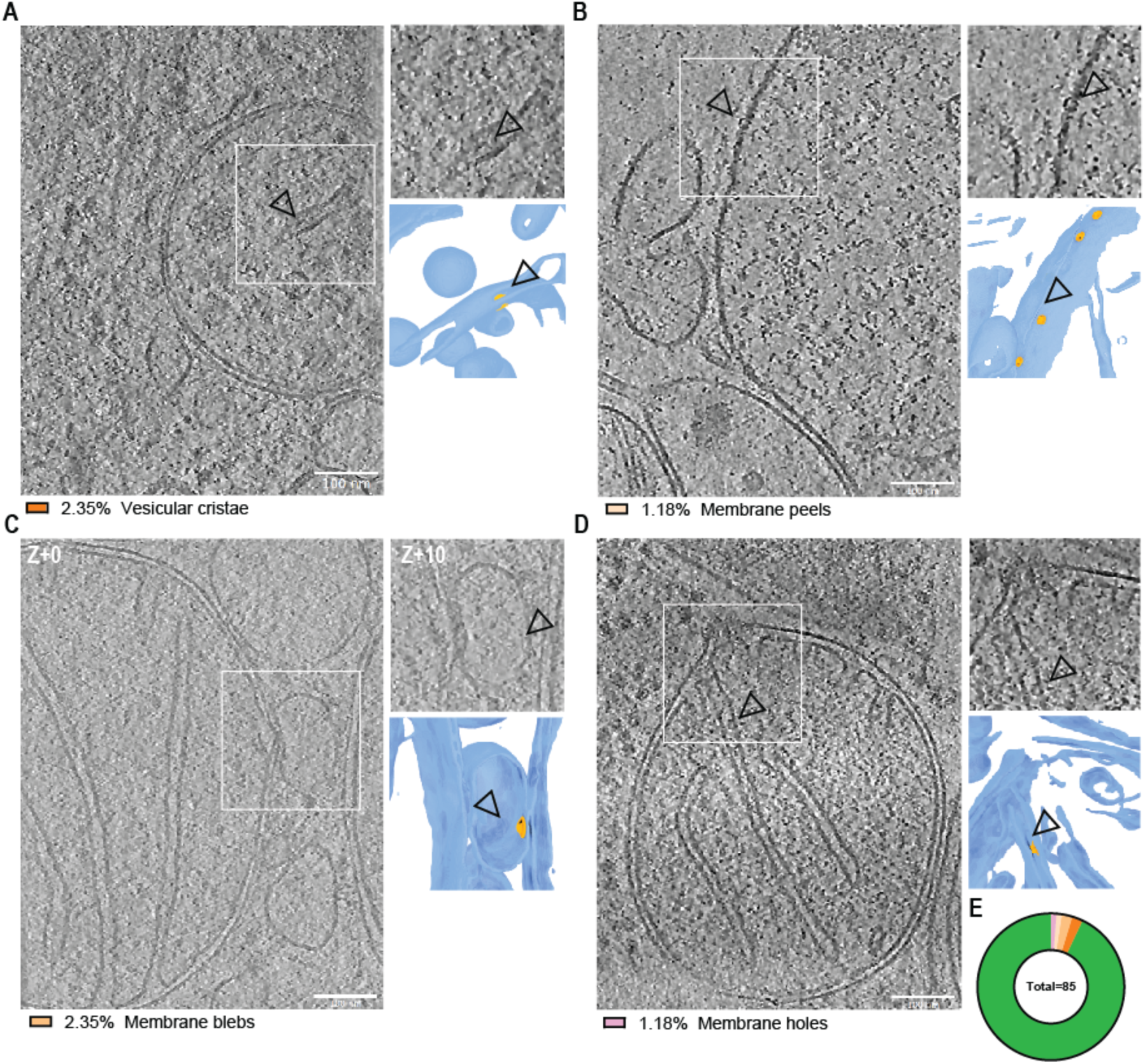
Prohibitin is an EM fiducial for studying mitochondrial membrane morphology changes. Back projection of prohibitin particles into raw tomograms identified mitochondria with membrane distortions. (A) A mitochondrial fragment with spherical cristae is juxtaposed by a single tubular crista containing prohibitin (inset). Prohibitin complexes were also identified exposed to the cytosol in mitochondria with outer membrane peels (B and inset). A single layer outer membrane bleb from a mitochondria is identified by a prohibitin complex on the interior of the bleb (C and inset). A double membrane rupture of a mitochondria is shown with prohibitin present in the cristae below the rupture site (D and inset). (E) Comparison of membrane morphologies from each class (n=85 fragments, 79 with no abnormalities).

**Supplementary Figure 5:**
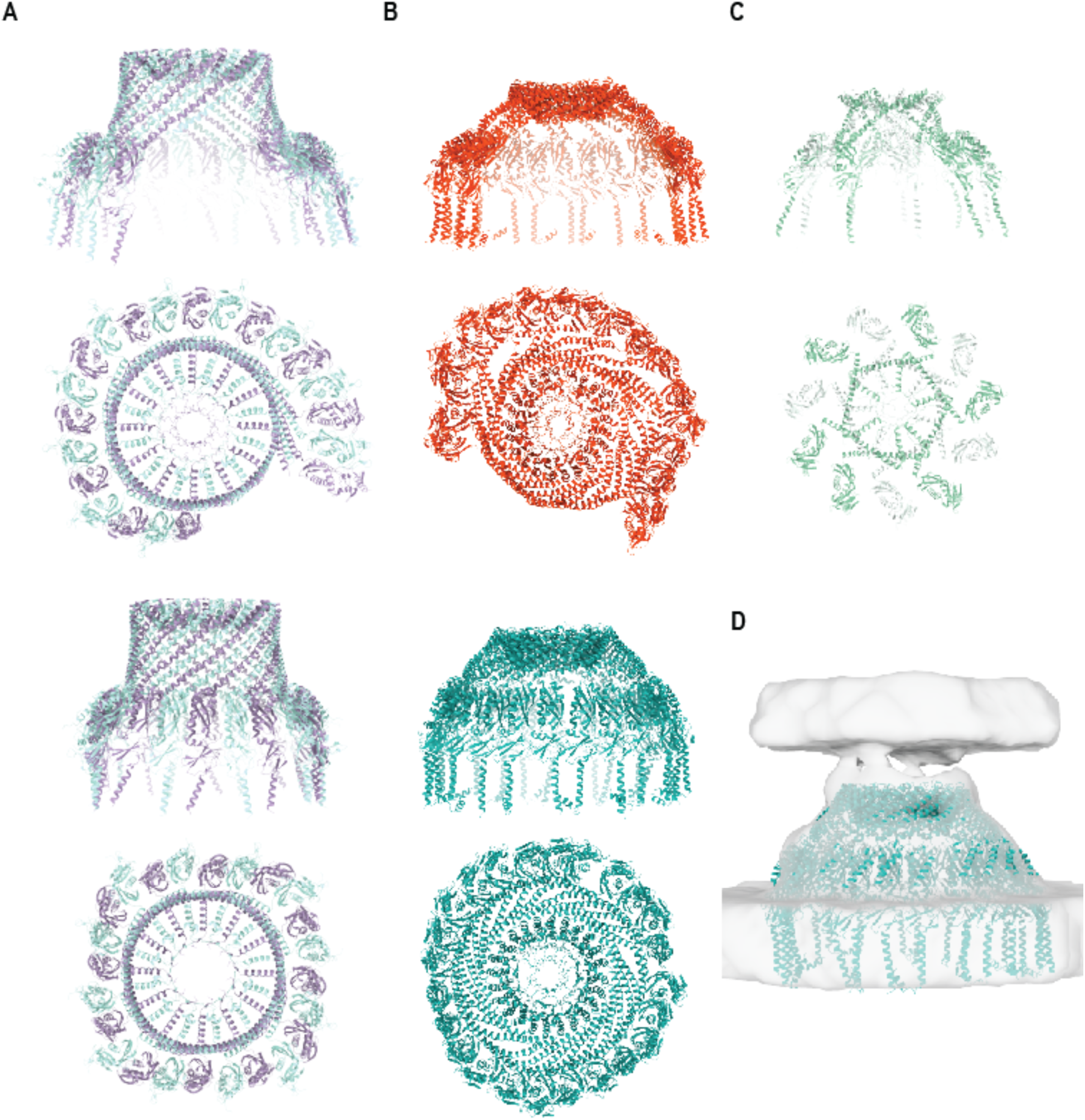
Structural comparison of HFLC/K and prohibitin structures. Structures of HFLC/K complex in the open (PDB: 9CZ2) and closed states (PDB: 7VHP) (A). Prohibitin models (open at top, closed at bottom) generated in this study (B). A prohibitin model generated from C11 symmetric processing (PDB: 8RRH) (C). Closed model from this study docked into the density map of putative prohibitin from Chlamydomonas (EMDB-50212) (D).

**Table S1.** Cryo-electron tomography data collection.

| Dataset # | 1 G3 | 2 G3 | 3 G4 | 4 G2 |
| --- | --- | --- | --- | --- |
| Grids | Quantifoil gold R2/2 | Quantifoil gold R2/2 | Quantifoil gold R2/2 | Quantifoil gold R2/2 |
| Cell type | U2OS | U2OS | U2OS | U2OS |
| Cryo-Specimen Freezing | Vitrobot Mark IV | Vitrobot Mark IV | Vitrobot Mark IV | Vitrobot Mark IV |
| Microscope, Voltage (keV) | Titan Krios G3, 300 | Titan Krios G3, 300 | Titan Krios G4, 300 | Titan Krios G3, 300 |
| Detector | Gatan Quantum K3 | Gatan Quantum K3 | Falcon i4 | Gatan Quantum K3 |
| Energy filter slit width (eV) | 25 | 25 | 10 | 20 |
| Electron Exposure ($e^-/\text{\AA}^2$) dose fractionation | ~90 | ~90 | ~120 | ~120 |
| Defocus Range ($\mu\text{m}$) | -2 $\mu\text{m}$ to -6 $\mu\text{m}$ | -2 $\mu\text{m}$ to -6 $\mu\text{m}$ | -2 $\mu\text{m}$ to -6 $\mu\text{m}$ | -2 $\mu\text{m}$ to -6 $\mu\text{m}$ |
| Tilt scheme | -60°/+60°, 3°, dose symmetrical (Hagen Scheme) | 60°/+60°, 3°, dose symmetrical (Hagen Scheme) | 60°/+60°, 3°, dose symmetrical (Hagen Scheme) | 60°/+60°, 3°, dose symmetrical (Hagen Scheme) |
| Movie recording | 6-8 | 6-8 | 9 | 6-8 |
| Magnification (times) | 43000 | 43000 | 64000 | 42000 |
| Pixel Size ($\text{\AA}/\text{px}$) | 1.05 (Super resolution) | 1.05 (Super resolution) | 1.965 | 0.90 (Super resolution) |
| Tomograms acquired | 21 | 50 | 47 | 33 |

## Movie titles

**Movie 1: Representative tomogram of an untreated mitochondria.**

**Movie 2: Representative tomogram of an OA treated mitochondria**

**Movie 3: Tomogram of a phagophore with putative BLTPs targeting a damaged mitochondrial fragment.**

**Movie 4: Tomogram showing membranes enveloping an OA treated mitochondrial fragment.**

**Movie 5: Morph of prohibitin maps and models illustrating differences between the two conformations.**

