## Supplementary material for "*In situ* cryo-ET visualization of mitochondrial depolarization and mitophagic engulfment": key resource table

| RESOURCE TYPE | RESOURCE NAME | SOURCE | IDENTIFIER | NEW/REUSE | ADDITIONAL INFORMATION |
|---|---|---|---|---|---|
| **Dataset** | structural coordinates | rcsb.org | in progress | NEW | |
| **Dataset** | density maps | ebi.ac.uk/emdb | in progress | NEW | |
| **Dataset** | raw cryo-EM data | ebi.ac.uk/empiar | in progress | NEW | |
| **Dataset** | | | | | |
| **Software/code** | Zeiss ZEN Blue software version 3.7 | Carl Zeiss LLC | RRID:SCR_013672 | REUSE | |
| **Software/code** | Arivis Vision4D (ver 4.2.1) | Carl Zeiss LLC | RRID:SCR_018000 | REUSE | |
| **Software/code** | Scipion | | RRID:SCR_016738 | REUSE | https://scipion.i2pc.es/ |
| **Software/code** | Membrain | | https://github.com/CellArchLab/MemBrain-v2 | REUSE | |
| **Software/code** | Dragonfly | | | REUSE | https://dx.doi.org/10.3791/64435-v |
| **Software/code** | Pytom model | | https://github.com/SBC-Utrecht/PyTom | REUSE | |
| **Software/code** | AlphaFold3 | | RRID:SCR_025885 | REUSE | |
| **Software/code** | ImageJ | | RRID:SCR_002285 | REUSE | https://imagej.net/ |
| **Software/code** | JACoP | | RRID: SCR_025164 | REUSE | https://imagej.net/plugins/jacop |
| **Software/code** | GraphPad Prism 9 | | RRID:SCR_002798 | REUSE | |
| **Software/code** | ISOLDE (ver 1.5) | PMID: 29872003 | n/a | REUSE | https://isolde.cimr.cam.ac.uk/ |
| **Software/code** | MotionCor3 | | https://github.com/czimaginginstitute/MotionCor3 | REUSE | |
| **Software/code** | 3DEM | | RRID:SCR_016738 | REUSE | https://github.com/3dem/relion |
| **Software/code** | CTFfind 5 | | https://github.com/GrigorieffLab/ctffind5_manuscript | REUSE | |
| **Software/code** | AreTomo2 | | https://github.com/czimaginginstitute/AreTomo2 | REUSE | |
| **Software/code** | ChimeraX (ver 1.5) | PMID: 32881101 | RRID:SCR_015872 | REUSE | https://www.cgl.ucsf.edu/chimera/ |
| **Software/code** | Napari | | RRID:SCR_022765 | REUSE | https://www.napari-hub.org/ |
| **Software/code** | Relion 5 | | https://github.com/3dem/relion | REUSE | |
| **Software/code** | GraphPad Prism (ver 10.1.0) | Graphstats Technologies | | REUSE | http://www.graphpad.com/ |
| **Software/code** | Watershed | | | REUSE | |
| **Protocols** | OA treatment and Quantification of Parkin recruitment to mitoch | protocols.io | https://dx.doi.org/10.17504/protocols.io.5qpvoox4dv4o/v1 | NEW | |
| **Protocols** | Transfection | protocols.io | http://dx.doi.org/10.17504/protocols.io.ewov1dr82vr2/v1 | NEW | |
| **Protocols** | Cell culture and cell line generation | protocols.io | https://dx.doi.org/10.17504/protocols.io.81wgbxq2qlpk/v1 | REUSE | |
| **Protocols** | Mitophagy flux assay via in-gel fluorescence | protocols.io | https://dx.doi.org/10.17504/protocols.io.x54v9pwkmg3e/v1 | REUSE | |
| **Protocols** | Lentivirus plasmids generation | protocols.io | https://dx.doi.org/10.17504/protocols.io.yxmvm3z5bl3p/v1 | REUSE | |
| **Protocols** | EM grid seeding and cryo-FIB milling | protocols.io | https://dx.doi.org/10.17504/protocols.io.dm6gpde65gzp/v1 | NEW | |
| **Protocols** | Cryo-electron tomography data acquisition | protocols.io | https://dx.doi.org/10.17504/protocols.io.36wgq6rkklk5/v1 | NEW | |
| **Protocols** | Cryo-electron tomography data processing and model building | protocols.io | https://dx.doi.org/10.17504/protocols.io.36wgq6rkklk5/v1 | NEW | |
| **Protocols** | Live Airyscan microscopy of mitochondrial ultrastructure | protocols.io | in progress | | |
| **Antibody** | / | | | | |
| **Bacterial strain** | / | | | | |
| **Virus strain** | / | | | | |
| **Biological sample** | / | | | | |
| **Chemical, peptide, or recombinant protein** | Lipofectamine | Thermo Fisher Scientific | REF:11668030 | REUSE | |
| **Chemical, peptide, or recombinant protein** | Opti-MEM | Thermo Fisher Scientific | REF: 31985-070 | REUSE | |
| **Critical commercial assay** | / | | | | |
| **Experimental model: Cell line** | U-2OS | Cell Culture Facility, UC Berkeley | RRID:SCR_017924 | REUSE | |
| **Experimental model: Cell line** | HEK293T | Cell Culture Facility, UC Berkeley | RRID:CVCL_0063 | REUSE | |
| **Oligonucleotide** | / | | | | |
| **Recombinant DNA** | mCherry-TOMM20-N-10 | AddGene | Plasmid # 55146 | REUSE | 500ng per dish |
| **Recombinant DNA** | ATP5F1B-tGFP | OriGene | NM_001686, SKU: RG201638 | NEW | 250ng per dish |
| **Recombinant DNA** | pCMV-VSV-G | Addgene | Addgene_8454 | REUSE | https://www.addgene.org/8454/ |
| **Recombinant DNA** | pCMVR8.74 | Addgene | Addgene_22036 | REUSE | https://www.addgene.org/22036/ |
| **Recombinant DNA** | pBS-CMV-gagpol | Addgene | Addgene_35614 | REUSE | https://www.addgene.org/35614/ |
| **Recombinant DNA** | pLV-mCherry-Parkin | Addgene | Addgene_237397 | NEW | |
| **Recombinant DNA** | pLV-BFP-mito | Addgene | Addgene_237398 | NEW | |
| **Recombinant DNA** | pMRX-IB-pSu9-HaloTag7-mGFP | Addgene | Addgene_184905 | REUSE | https://www.addgene.org/184905/ |
| **Recombinant DNA** | pMK1253 | Addgene | Addgene_133058 | REUSE | https://www.addgene.org/133058/ |
| **Recombinant DNA** | pBMN-mCherry-Parkin | Addgene | RRID:Addgene_59419 | REUSE | https://www.addgene.org/59419/ |
| **Recombinant DNA** | EBFP2-Mito-7 | Addgene | Addgene_55248 | REUSE | https://www.addgene.org/55248/ |
